# PRDM14 controls X-chromosomal and global epigenetic reprogramming of H3K27me3 in migrating mouse primordial germ cells

**DOI:** 10.1101/551895

**Authors:** Anna Mallol, Maria Guirola, Bernhard Payer

## Abstract

In order to prepare the genome for gametogenesis, primordial germ cells (PGCs) undergo extensive epigenetic reprogramming during migration towards the gonads in mammalian embryos. This includes changes on a genome-wide scale and additionally in females the remodeling of the inactive X-chromosome to enable X-chromosome reactivation (XCR). However, if global and X-chromosomal remodeling are related and which factors are important is unknown. Here we identify the germ cell determinant PR-domain containing protein 14 (PRDM14) as the first known factor that is instrumental for both global and X-chromosomal reprogramming in migrating mouse PGCs. We find that global upregulation of the repressive histone H3 lysine 27 trimethylation (H3K27me3) mark is PRDM14 dosage-dependent in PGCs of both sexes. When focusing on XCR, we observed that PRDM14 is required for removal of H3K27me3 from the inactive X-chromosome. Furthermore we show that global and X-chromosomal H3K27me3 reprogramming are functionally separable, despite their common regulation by PRDM14. Thereby we provide mechanistic insight and spatiotemporal resolution to the remodeling of the epigenome during mouse PGC migration and link epigenetic reprogramming to its developmental context *in vivo*.

## INTRODUCTION

The germ cell lineage has the unique function of transmitting genetic and epigenetic information from one generation to the next. Mammalian germ cell development is an intricate process, which involves extensive changes to the epigenome before it is passed on by the gametes. In mice, germ cell development begins with specification of PGCs in the proximal epiblast. After specification, PGCs migrate through the hindgut in order to reach the genital ridges (future gonads), where they later undergo meiosis and sex-specific differentiation into eggs and sperm. During migration and colonialization of the gonads, the PGC epigenome is remodeled at multiple levels, which includes global DNA-demethylation, erasure of genomic imprints, removal of histone H3 lysine 9 dimethylation (H3K9me2) and a global increase of the PRC2 (Polycomb repressive complex 2)-associated H3K27me3 mark (Hajkova et al., 2008; Leitch et al., 2013a; Reik and Surani, 2015; Saitou et al., 2012; Seki et al., 2005; Seki et al., 2007). Around the same time, in female PGCs, the inactive X-chromosome is reactivated by XCR, which involves downregulation of the non-coding RNA Xist, the master regulator of X-inactivation, removal of repressive epigenetic marks like H3K27me3 and reactivation of X-linked genes (Chuva de Sousa Lopes et al., 2008; de Napoles et al., 2007; Payer, 2016; Sugimoto and Abe, 2007). Interestingly, H3K27me3 is therefore removed from the inactive X-chromosome while it increases globally on autosomes. It is currently unknown, how X-chromosomal and global H3K27me3 reprogramming occur along the migration path of PGCs and if they are functionally linked or are rather regulated independently of each other.

XCR is a process linked to naïve pluripotency and germ cell fate (Pasque and Plath, 2015; Payer and Lee, 2014). While XCR-kinetics have been characterized during mouse and human germ cell development (Chuva de Sousa Lopes et al., 2008; de Napoles et al., 2007; Guo et al., 2015; Sugimoto and Abe, 2007; Tang et al., 2015), the molecular mechanisms are largely unknown. Although many factors have been described, which are instrumental in setting up the inactive chromatin configuration during X-chromosome inactivation (Galupa and Heard, 2018; Moindrot and Brockdorff, 2016; Payer, 2017), little is known about the factors, which are functionally important to reverse the inactive state during XCR. Identifying such X-reactivation factors would be a key step towards revealing mechanistically how epigenetic memory can be erased in the germ line. We and others have shown that the transcriptional regulator PRDM14 plays a critical role in XCR in the mouse blastocyst epiblast *in vivo* and during induced pluripotent stem cell (iPSC) reprogramming (Payer et al., 2013) and epiblast stem cell (EpiSC) to embryonic stem cell (ESC) conversion (Gillich et al., 2012) *in vitro*. In the absence of PRDM14, downregulation of Xist and removal of the H3K27me3 mark from the inactive X-chromosome are perturbed, likely due to repressive functions of PRDM14 on *Xist* and its activator *Rnf12/Rlim* (Ma et al., 2011; Payer et al., 2013).

Besides being important for naïve pluripotency (Yamaji et al., 2013) and the associated XCR, PRDM14 is also critical for germ cell development. *Prdm14*-mutant mouse embryos show reduced PGC numbers (Yamaji et al., 2008), and overexpression of PRDM14 in epiblast-like cells (EpiLCs) is sufficient to induce germ cell fate *in vitro* (Nakaki et al., 2013a). Importantly, PRDM14 is a regulator of global epigenetic changes both in pluripotent stem cells as well as in PGCs (Nakaki et al., 2013b; Seki, 2018). In particular, it has been shown that PRDM14 is responsible for the low global DNA-methylation levels characteristic of naïve pluripotent stem cells and PGCs, both by repressing DNA-methyltransferase genes, as well as by recruiting DNA-demethylases of the Ten-eleven translocation (TET) family (Ficz et al., 2013; Grabole et al., 2013; Leitch et al., 2013b; Okashita et al., 2014; Yamaji et al., 2008; Yamaji et al., 2013). Besides controlling DNA-hypomethylation, PRDM14 is also important for global remodeling of histone marks in pluripotent stem cells and PGCs. During mouse preimplantation development, PRDM14 shows asymmetric expression in blastomeres of the 4-cell embryo and has been implicated in driving cells towards inner cell mass fate by recruiting CARM1, a histone H3 arginine 26 (H3R26me2) methyltransferase (Burton et al., 2013). In migrating PGCs, *Prdm14*-mutant mouse embryos fail to downregulate the repressive H3K9me2 mark and its associated histone methyltransferase GLP/EHMT1 (Yamaji et al., 2008), most likely because *Glp/Ehmt1* is a directly repressed target gene of PRDM14 (Tu et al., 2016). The PGC-specific upregulation of H3K27me3 on a global level is also impaired in *Prdm14-/-* embryos (Yamaji et al., 2008) and genome-wide distribution of H3K27me3 is misregulated in *Prdm14-/-* ESCs (Yamaji et al., 2013).

Despite the known importance of PRDM14 for epigenetic remodeling in the germ cell lineage, how PRDM14 controls its different aspects *in vivo* is largely unresolved: For example, it remains unknown, if PRDM14 is important for X-chromosome reactivation also in female PGCs, and if this is coordinated with genome-wide epigenetic changes occurring at that time. Furthermore no information is available about the spatial distribution of germ cells along their migration path towards the gonads during epigenetic reprogramming. To address these questions, we investigated in this study epigenetic remodeling of the polycomb-associated H3K27me3 mark and its dependency on PRDM14 in migrating mouse PGCs. By using a whole-mount embryo staining approach, we were able for the first time to collect spatiotemporal information about the epigenetic reprogramming process in mouse PGCs in relation to their migratory progress. We found that PRDM14 is critical both for X-chromosomal removal as well as global upregulation of H3K27me3 in PGCs. We furthermore find differences in PRDM14-dosage dependence and kinetics of X-chromosomal and global H3K27me3 changes. While both events are controlled by PRDM14, they do not depend on each other and therefore must be regulated through different mechanisms. In summary, we describe here how PRDM14 regulates key aspects of epigenetic changes in germ cells and how this is linked with progression of germ cell migration and development.

## RESULTS AND DISCUSSION

### PRDM14 controls number and local distribution of PGCs during their migration

To address the function of PRDM14 for PGC number and migration, we immunostained whole mount embryonic day (E)9.5 mouse embryos from *Prdm14^+/−^* x *Prdm14^+/−^* heterozygous crosses for AP2γ, a specific marker and critical factor for PGC development (Nakaki et al., 2013a; Weber et al., 2010). We found that migrating PGCs at this stage were spread throughout the hindgut, and observed in *Prdm14^-/-^* embryos a strong depletion of PGCs or even their absence (7/17 embryos), when compared to wildtype and heterozygous littermates (Fig. 1A, S1, Table S1), which is consistent with a previous study of this mouse line (Yamaji et al., 2008). Generally, PGC numbers increased with developmental progression (somite number) in *Prdm14^+/+^* and *Prdm14^+/−^* embryos, while it remained low in *Prdm14^-/-^* embryos (Fig. 1B). This is not due to a general developmental delay of *Prdm14*-mutant embryos at E9.5, as somite numbers were similar between different *Prdm14* genotypes (Fig. S2A, Table S1).

**Fig. 1.**
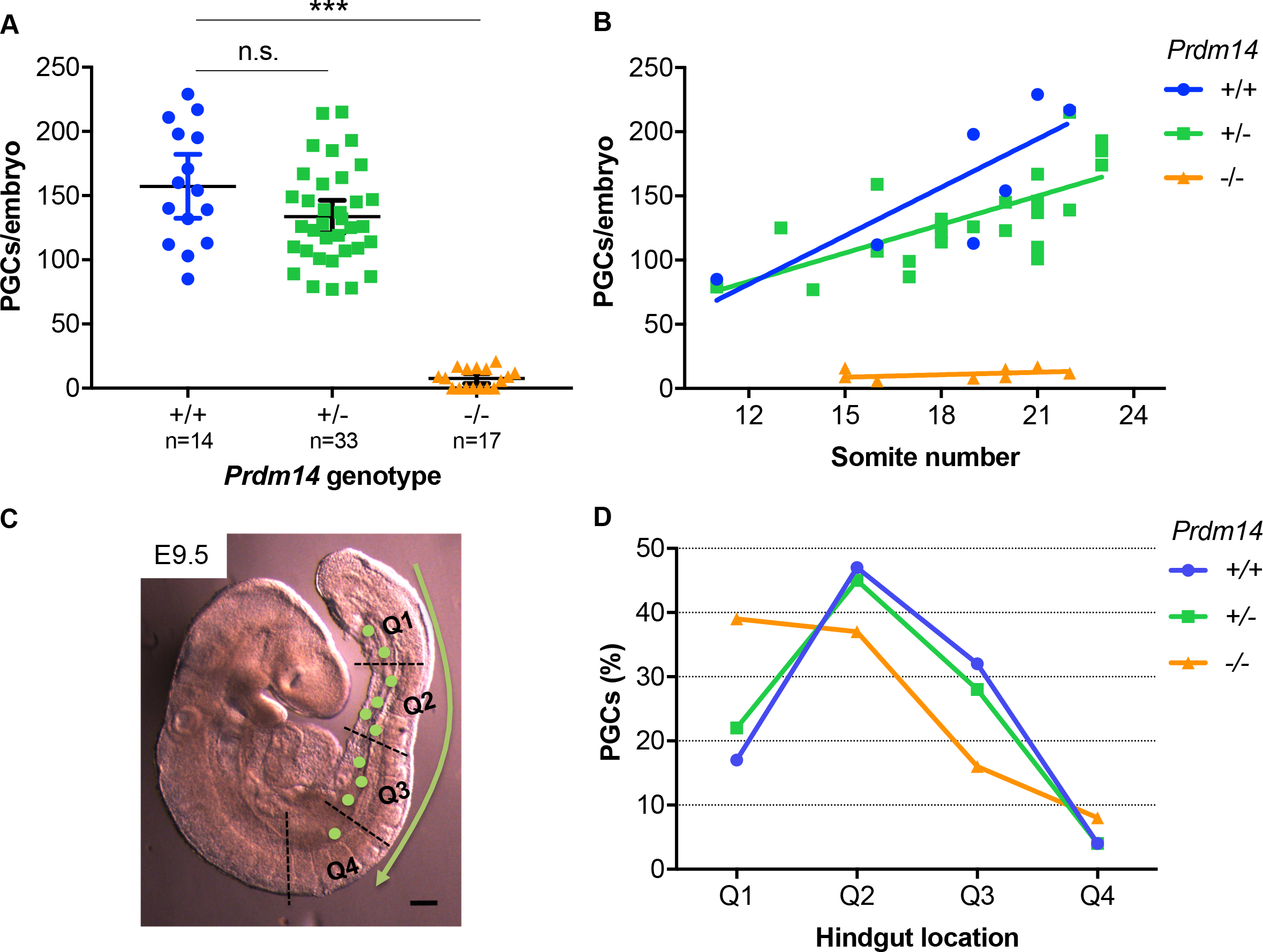
*Prdm14^-/-^* E9.5 embryos have reduced PGC numbers with altered distribution along the hindgut. (A) Numbers of AP2γ-positive PGCs per embryo across *Prdm14* genotypes. Means and confidence intervals are shown. Kruskal-Wallis test was used for statistical comparison (p<0.001). (B) Linear regression of AP2γ-positive PGC-numbers versus developmental stage (somite number). R square values for *Prdm14* +/+, +/− and -/- embryos were 0.659, 0.514 and 0.001, respectively. (C) Image of a E9.5 mouse embryo with PGCs (drawn green dots) migrating along the hindgut divided into four quadrants (Q1 – Q4). Scale bar: 250 μm. (D) Distribution of PGCs along the hindgut across *Prdm14* genotypes.

In order to analyze the distribution of PGCs in more detail, we divided the migration path along the hindgut into 4 quadrants (Fig. 1C); starting with quadrant 1 (Q1) at the posterior end at the base of the allantois and finishing with quadrant 4 (Q4) near the area where PGCs would exit the hindgut and enter the genital ridges (future gonads). At E9.5, PGCs would be found mostly in Q2 and Q3 in *Prdm14^+/+^* and *Prdm14^+/−^* embryos, while PGCs in *Prdm14^-/-^* embryos would be located predominantly in Q1 and Q2 (Fig. 1D, S2B). This indicates that *Prdm14*-mutant PGCs might be delayed in migration, although few of them can be also found in Q3 and Q4. In summary, *Prdm14-*mutant embryos show a strong reduction in PGC-numbers and a defect in PGC migration.

### PRDM14 is required for global upregulation of H3K27me3 in migrating PGCs

As PRDM14 has a role in the global epigenetic reprogramming occurring in PGCs (Yamaji et al., 2008), we wanted to investigate in more detail its function for upregulating the H3K27me3 mark, which is a hallmark of migrating PGCs (Hajkova et al., 2008; Seki et al., 2005; Seki et al., 2007). We therefore costained E9.5 embryos of different *Prdm14* genotypes with AP2γ and H3K27me3 antibodies (Fig. 2A). We scored H3K27me3 as being upregulated or non-upregulated in AP2γ-positive PGCs when compared to surrounding somatic cells. It thereby became apparent that H3K27me3 upregulation was in direct relation to PRDM14 dosage, as about 78% of wildtype PGCs had elevated H3K27me3 staining, while only 54% of heterozygote and only 19% of Prdm14-null PGCs did (Fig. 2B, Table S1). These differences were observed to a similar degree both in male and in female embryos, indicating that PRDM14 controlled global H3K27me3 levels in a sex-independent manner.

**Fig. 2.**
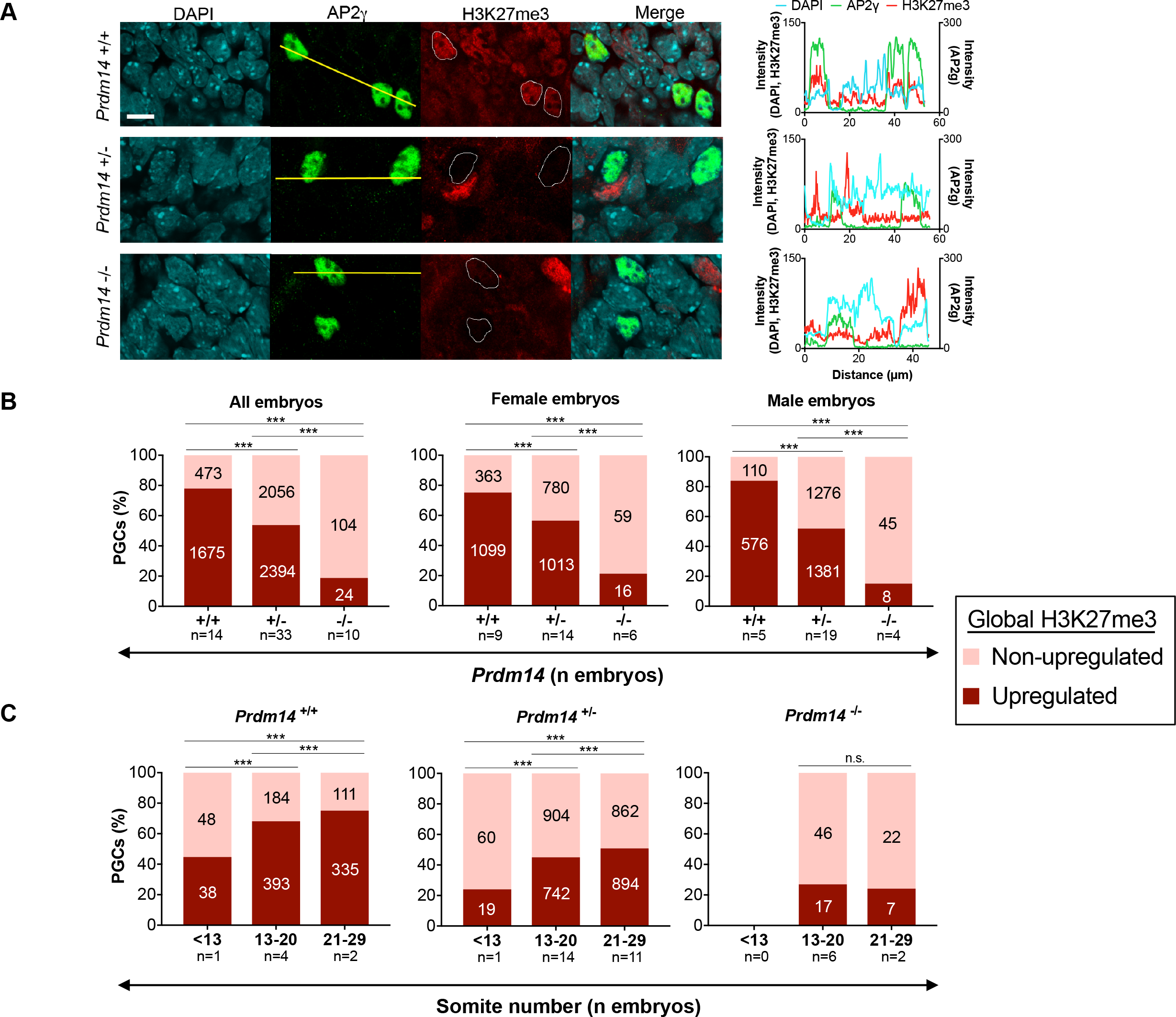
PRDM14 dosage controls global H3K27me3-levels in migrating PGCs in both sexes. (A) Representative images of AP2γ-positive PGCs with upregulated and non-upregulated global levels of H3K27me3 across different *Prdm14* genotypes (PGC-nuclei are outlined in H3K27me3 images). Scale bar: 10 μm. Intensity values for each channel along the yellow lines are plotted on the right. (B) Percentages of H3K27me3-upregulated and non-upregulated PGCs across *Prdm14* genotypes in male and female embryos. (C) Percentages of H3K27me3 upregulated and non-upregulated PGCs at different developmental stages (somite number). Labels in each column indicate number of PGCs analyzed. Chi-square test was used for statistical comparisons (p<0.001).

When we assessed H3K27me3 upregulation in relation to developmental progression (somite number) at E9.5, we observed that H3K27me3 increased with somite number in *Prdm14^+/+^* and *Prdm14^+/−^*, but not in *Prdm14^-/-^* embryos (Fig. 2C). Interestingly, H3K27me3 levels did not seem to vastly change in PGCs within different quadrants of their migration path, indicating that global H3K27me3 upregulation depends more on overall developmental progression of the embryo than on the position of the PGCs along the hindgut (Fig. S3). Overall, we conclude that global H3K27me3 upregulation in PGCs is dependent on PRDM14 dosage and progresses with developmental stage at equal rate in both sexes.

### X-chromosomal H3K27me3-erasure during XCR in female PGCs is dependent on PRDM14

In contrast to the global increase of H3K27me3, stands the erasure of H3K27me3 from the inactive X-chromosome in female PGCs during the process of XCR (Chuva de Sousa Lopes et al., 2008; de Napoles et al., 2007). As these events occur around the same time and as XCR is controlled by PRDM14 in mouse blastocysts and during iPSC reprogramming (Payer et al., 2013), we wanted to know if PRDM14 is also required for XCR in PGCs. Thus we scored the disappearance of the distinctive H3K27me3 accumulation from the inactive X-chromosome (“H3K27me3-spot”) in female PGCs in embryos of different *Prdm14* genotypes (Fig. 3A). While about half of both *Prdm14* wildtype (52%) and heterozygous (51%) PGCs have erased the H3K27me3-spot from the X, only around 23% of *Prdm14*-mutant PGCs have lost the spot at E9.5 (Fig. 3B, Table S1). The erasure of the H3K27me3 spot occurred progressively in *Prdm14^+/+^* and *Prdm14^+/−^* PGCs during their migration along the hindgut, but did not increase with migration in *Prdm14^-/-^* PGCs (Fig. 3C). In contrast to global H3K27me3 upregulation, removal of the X-chromosomal H3K27me3-spot therefore did not seem to be PRDM14-dose dependent and did not change substantially between E9.5 embryos of different somite number (Fig. S4). Taken together, our results show that similar to its role in the pluripotent epiblast and during iPSC reprogramming (Payer et al., 2013), PRDM14 is also a key factor for XCR in PGCs by promoting the loss of the X-chromosomal H3K27me3 mark.

**Fig. 3.**
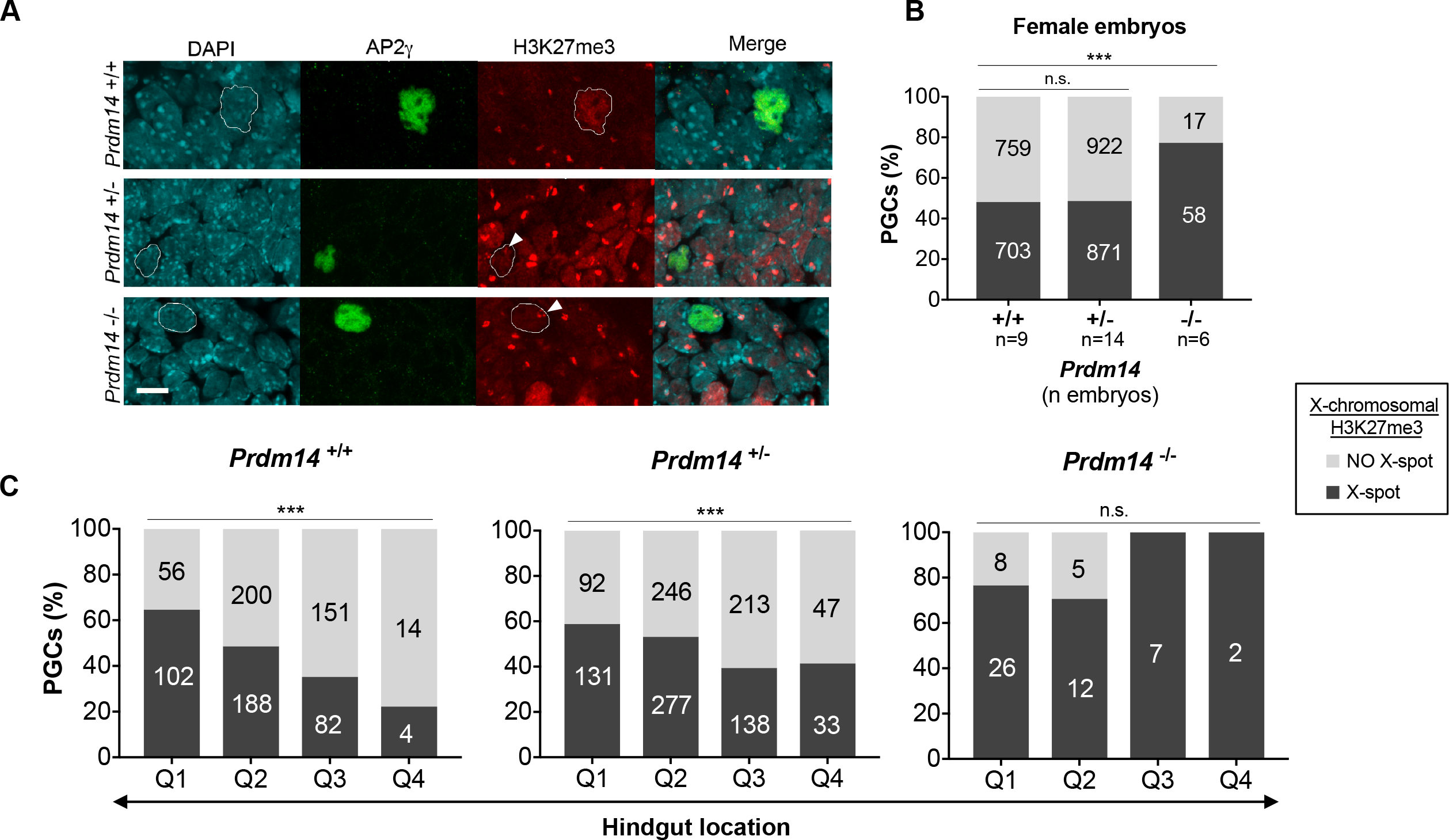
Erasure of H3K27me3-enrichment from the inactive X-chromosome in migrating female PGCs is dependent on PRDM14. (A) Representative images of AP2γ-positive PGCs with presence or absence of H3K27me3-accumulation (H3K27me3 spot, white arrowheads) on the inactive X-chromosome. PGC-nuclei are outlined in the H3K27me3 channel. Scale bar: 10 *μ*m. (B) Percentage of H3K27me3 spot-positive and -negative PGCs across *Prdm14* genotypes. Labels in each column indicate number of cells analyzed. (C) Erasure of the H3K27me3-spot in PGCs of different *Prdm14* genotypes during their migration along the hindgut divided in four quadrants (Q1-Q4). Labels in each column indicate number of PGCs analyzed. Chi-square test was used for statistical comparisons (p<0.001).

### X-chromosomal and global H3K27me3 reprogramming occur independently in female PGCs

H3K27me3 and its associated enzymatic complex, PRC2, are strongly enriched on the inactive X-chromosome in female cells. Therefore, we hypothesized, if this could act as a “sink” for PRC2, whereby global upregulation of H3K27me3 in PGCs would require first stripping PRC2 off the X-chromosome through XCR to make it available for acting elsewhere in the genome. In order to test this hypothesis, we investigated the relationship between X-chromosomal downregulation and global upregulation of H3K27me3 in female PGCs (Fig. 4, S5). If the inactive X were acting as a sink for PRC2, we would expect global H3K27me3 not to be upregulated in PGCs harboring an X-spot. However, when comparing within the different genotypes the fraction of PGCs which have lost or retained the X-chromosomal H3K27me3-spot, we did not detect any significant differences in global H3K27me3 upregulation (Fig. 4A). This indicates that losing the X-spot is not a general prerequisite for global H3K27me3 upregulation in female PGCs. Also the fact that *Prdm14^+/+^* and *Prdm14^+/−^* PGCs showed almost identical X-spot loss (52% in +/+ vs. 51% in +/−) but different global H3K27me3 upregulation levels (75% in +/+ vs. 56% in +/−), indicates that X-spot loss and global H3K27me3 upregulation are independent epigenetic events. In conclusion, while PRDM14 is both required for normal X-chromosomal and global H3K27me3 reprogramming in PGCs (Fig. 4B, S6), these events are mechanistically separable and do not depend on each other.

**Fig. 4.**
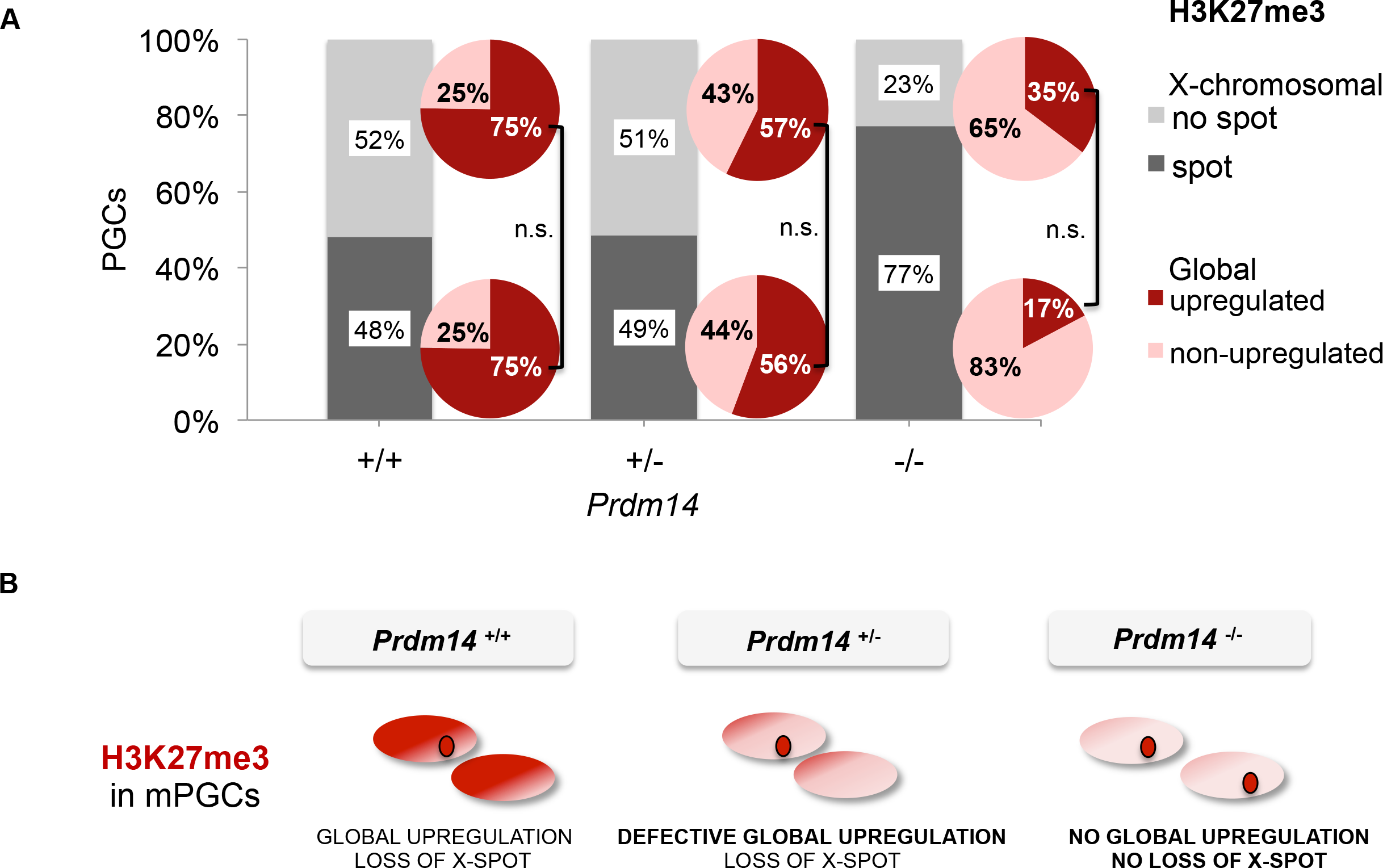
Changes in both X-chromosomal and global levels of H3K27me3 in female PGCs require PRDM14 but are independent of each other. (A) Relationship between X-chromosomal (bar graphs, % of PGCs per genotype) and global (pie charts, % of PGCs per genotype within each H3K27me3 spot/no spot sub-category) levels of H3K27me3 in PGCs from female embryos of different *Prdm14* genotypes. Chi-square test was used to compare dependency of global of H3K27me3 upregulation on X-chromosomal removal for each *Prdm14* genotype (n.s., not significant; *Prdm14^+/+^* p=0.956, *Prdm14^+/−^* p=0.499, *Prdm14^-/-^* p=0.110). (B) Schematic summary of *Prdm14* genotype effect on global and X-chromosomal H3K27me3 reprogramming in mouse PGCs.

### Summary and discussion

In this study, we have investigated the role of PRDM14 during epigenetic reprogramming in migrating mouse PGCs (summarized in Fig. 4B, S6). We thereby have uncovered multiple roles for PRDM14 during germ cell development. Most importantly, we have identified PRDM14 to our knowledge as the first factor with functional importance for XCR in the germ cell lineage. In particular, we found that erasure of H3K27me3 mark from the inactive X-chromosome, a key step during X-reactivation (Borensztein et al., 2017; Chuva de Sousa Lopes et al., 2008; de Napoles et al., 2007; Mak et al., 2004), occurred progressively during the migration of female PGCs and required PRDM14. This observation is in line with the role of PRDM14 during X-reactivation in the mouse blastocyst and during iPSC-reprogramming (Payer et al., 2013).

Second, we uncovered that global H3K27me3 upregulation in PGCs is dependent on PRDM14 in a dosage-sensitive manner, as *Prdm14^+/−^* and *Prdm14^-/-^* PGCs showed a progressive defect. Interestingly, we observed that H3K27me3 upregulation seemed to mostly depend on the developmental stage of the embryo rather than the PGC position along the migratory path, which is in contrast to the kinetics of X-specific H3K27me3 removal.

Another key finding from our study is that the global upregulation and X-chromosomal depletion of the H3K27me3 mark seem to be controlled by distinct mechanisms despite their common dependency on PRDM14. This is based on several observations: First, global H3K27me3 upregulation is sensitive to PRDM14 dosage, while X-specific H3K27me3 removal is not. Second, global upregulation correlates with embryonic stage (somite number), while X-chromosomal H3K27me3 loss occurs progressively during PGC migration. Third, global H3K27me3 upregulation occurs with similar kinetics in male and female embryos and is affected to a similar extent by loss of PRDM14 in both sexes. Finally, X-chromosomal depletion of H3K27me3 does not seem to be required for global H3K27me3 upregulation on a single-cell level. How then could PRDM14 control H3K27me3 remodeling on autosomes versus on the X-chromosome? On the global scale, previous studies suggested that PRDM14 could directly interact with the PRC2 complex in pluripotent stem cells and thereby facilitate its recruitment to PRDM14 target genes (Chan et al., 2013; Yamaji et al., 2013) or that PRDM14 could activate expression of the PRC2 component *Suz12* (Grabole et al., 2013). More recent studies however challenged this view and proposed that PRDM14 mainly acts though its binding partner and co-repressor CBFA2T2/MTGR1 (Kawaguchi et al., 2019; Nady et al., 2015; Tu et al., 2016), suggesting that PRC2 might be recruited secondarily to PRDM14 targets. Regarding its X-chromosome-specific role, we have previously shown that PRDM14 binds to regulatory regions at *Xist* intron 1 and upstream of the *Xist*-activator *Rnf12* (Payer et al., 2013) and thereby facilitates repression of *Xist* during XCR. As H3K27me3-accumulation on the X-chromosome is *Xist*-dependent (Plath et al., 2003; Schoeftner et al., 2006; Silva et al., 2003; Zhao et al., 2008), PRDM14 might thereby regulate X-chromosomal H3K27me3-depletion by downregulating *Xist*. The fact that H3K27me3-removal from the X-chromosome occurs progressively in PGCs along their migration path, could speak in favor of a cell-proliferation and replication-dependent, passive dilution mechanism, similar as it has been proposed for global DNA-demethylation in PGCs (Kagiwada et al., 2013; Seisenberger et al., 2012). Nevertheless, also active mechanisms could play a role like removal of the mark by the H3K27me3-demethylase KDM6A/UTX, which has a partial effect on H3K27me3-demethylation during XCR in mouse blastocysts (Borensztein et al., 2017).

Finally, we have found that *Prdm14^-/-^* embryos displayed severely compromised germ cell numbers as reported previously (Yamaji et al., 2008) and a delay in PGC migration. This is in line with a recent study (Cheetham et al., 2018), which found that genes implicated in cell migration are bound by PRDM14 in PGC-like cells derived *in vitro*, suggesting that PRDM14 might be directly involved in controlling PGC migration.

In summary, here we have shown that autosomal and X-chromosomal reprogramming of H3K27me3 occur independently during PGC migration, but that they rely both on the key germ cell factor PRDM14. By adding to our understanding of the mechanisms underlying epigenetic changes required for germ cell development *in vivo*, we provide a framework for assessing and improving the quality of *in vitro*-derived gametes.

## MATERIALS AND METHODS

### Embryo isolation

Mouse care and procedures were conducted according to the protocols approved by the Ethics Committee on Animal Research of the Parc de Recerca Biomèdica de Barcelona (PRBB) and by the Departament de Territori i Sostenibilitat of the Generalitat de Catalunya.

*Prdm14* mutant mice (Yamaji et al., 2008) were maintained in a predominant C57BL/6 strain background. *Prdm14* heterozygous mice were mated and resulting embryos were harvested from pregnant females at E9.5 and dissected from maternal tissues.

### Whole mount immunostaining

The whole mount embryo immunostaining was performed as described in (Yamaji et al., 2008). The primary antibodies used were rabbit polyclonal anti-AP2γ (TCFAP2C) (Santa Cruz Biotechnology sc-8977) and mouse monoclonal anti-H3K27me3 (Active Motif 61707, clone MABI0323). The secondary antibodies used were donkey anti-rabbit IgG Alexa Fluor 488 and goat anti-mouse IgG Alexa Fluor 555 (Molecular Probes A21206 and A21424).

After the immunostaining, the somite number of every embryo was counted under the stereomicroscope. Then the embryo was split in two parts: the head was used for genotyping and the hindgut was dissected and mounted in Vectashield (Vector Laboratories) for observation with confocal microscopy.

### Image capture and analysis

Bright field images of the whole embryos were captured on a stereomicroscope using the Leica application suite software (Leica Microsystems, Wetzlar, Germany). Fluorescence imaging of the hindguts was performed on an inverted Leica TCS SP5 confocal microscope using the Leica Application Suite Advanced Fluorescence software. Z-stack images (1.5 μm intervals) of the full hindgut were acquired. Color setting and image processing were performed in Fiji (Schindelin et al., 2012) and Volocity (PerkinElmer) software.

The location in the hindgut and both the H3K27me3 global levels and the presence of the X-spot of every AP2γ positive cell were recorded. The global nuclear intensity of H3K27me3 staining was classified as “upregulated” or “non-upregulated” after comparing it with the staining intensity of the surrounding somatic cells in the same focus plane. The total length of the hindgut was divided in four quadrants (Q1-Q4 from the tail tip to the genital ridges, Figure 1C) to ease the analysis of the location of AP2γ positive cells.

### *Prdm14* and sex genotyping

*Prdm14* genotype was determined by polymerase chain reaction after image analysis. Primer sequences are detailed in (Yamaji et al., 2008).

The sex of the embryo was determined by the presence of the H3K27me3 spot corresponding to the silent X chromosome in somatic cells of female embryos (Plath et al., 2003).

### Statistical analysis

Several E9.5 litters were collected and the results from all embryos were pooled and analyzed with IBM SPSS Statistics software. For qualitative variables χ^2^ test was used, and for quantitative variables the non-parametric Kruskal–Wallis test was used. Pairwise comparisons with Bonferroni’s correction were performed using the Mann–Whitney U test. For lineal regression modeling, R^2^ coefficient was calculated. In all cases, p<0.05 was considered statistically significant.

## Acknowledgements

We thank members of the Payer lab for discussions and suggestions. We thank Mitinori Saitou for sharing the *Prdm14-KO* mouse strain and for advice on whole mount detection of PGCs by immunostaining. We also thank Guillaume Filion for critical reading and advice on statistical analysis. We thank the PRBB Animal Facility for mouse husbandry, the CRG Media kitchen for reagent preparation and the CRG Advanced Light Microscopy Unit for imaging support.

## Author contributions

Conceptualization and project design: A.M., M.G., B.P.; Experimental execution: A.M., M.G.; Data Collection: A.M., M.G.; Statistical Analysis A.M.; Writing: A.M., P.B.; Supervision + Funding Acquisition: B.P.

## References

Borensztein, M., Okamoto, I., Syx, L., Guilbaud, G., Picard, C., Ancelin, K., Galupa, R., Diabangouaya, P., Servant, N., Barillot, E., et al. (2017). Contribution of epigenetic landscapes and transcription factors to X-chromosome reactivation in the inner cell mass. Nat. Commun. 8, 1297.

Burton, A., Muller, J., Tu, S., Padilla-Longoria, P., Guccione, E. and Torres-Padilla, M.-E. (2013). Single-cell profiling of epigenetic modifiers identifies PRDM14 as an inducer of cell fate in the mammalian embryo. Cell Rep. 5, 687–701.

Chan, Y.-S., Göke, J., Lu, X., Venkatesan, N., Feng, B., Su, I.-H. and Ng, H.-H. (2013). A PRC2-dependent repressive role of PRDM14 in human embryonic stem cells and induced pluripotent stem cell reprogramming. Stem Cells 31, 682–92.

Cheetham, S. W., Gruhn, W. H., van den Ameele, J., Krautz, R., Southall, T. D., Kobayashi, T., Surani, M. A. and Brand, A. H. (2018). Targeted DamID reveals differential binding of mammalian pluripotency factors. Development 145, dev170209.

Chuva de Sousa Lopes, S. M., Hayashi, K., Shovlin, T. C., Mifsud, W., Surani, M. A. and McLaren, A. (2008). X chromosome activity in mouse XX primordial germ cells. PLoS Genet. 4, e30.

de Napoles, M., Nesterova, T. and Brockdorff, N. (2007). Early loss of Xist RNA expression and inactive X chromosome associated chromatin modification in developing primordial germ cells. PLoS One 2, e860.

Ficz, G., Hore, T. A., Santos, F., Lee, H. J., Dean, W., Arand, J., Krueger, F., Oxley, D., Paul, Y.-L., Walter, J., et al. (2013). FGF signaling inhibition in ESCs drives rapid genome-wide demethylation to the epigenetic ground state of pluripotency. Cell Stem Cell 13, 351–9.

Galupa, R. and Heard, E. (2018). X-Chromosome Inactivation: A Crossroads Between Chromosome Architecture and Gene Regulation. Annu. Rev. Genet. 52, 535–566.

Gillich, A., Bao, S., Grabole, N., Hayashi, K., Trotter, M. W. B., Pasque, V., Magnúsdóttir, E. and Surani, M. A. (2012). Epiblast stem cell-based system reveals reprogramming synergy of germline factors. Cell Stem Cell 10, 425–39.

Grabole, N., Tischler, J., Hackett, J. A., Kim, S., Tang, F., Leitch, H. G., Magnúsdóttir, E. and Surani, M. A. (2013). Prdm14 promotes germline fate and naive pluripotency by repressing FGF signalling and DNA methylation. EMBO Rep. 14, 629–37.

Guo, F., Yan, L., Guo, H., Li, L., Hu, B., Zhao, Y., Yong, J., Hu, Y., Wang, X., Wei, Y., et al. (2015). The Transcriptome and DNA Methylome Landscapes of Human Primordial Germ Cells. Cell 161, 1437–1452.

Hajkova, P., Ancelin, K., Waldmann, T., Lacoste, N., Lange, U. C., Cesari, F., Lee, C., Almouzni, G., Schneider, R. and Surani, M. A. (2008). Chromatin dynamics during epigenetic reprogramming in the mouse germ line. Nature 452, 877–881.

Kagiwada, S., Kurimoto, K., Hirota, T., Yamaji, M. and Saitou, M. (2013). Replication-coupled passive DNA demethylation for the erasure of genome imprints in mice. EMBO J. 32, 340–53.

Kawaguchi, M., Sugiyama, K., Matsubara, K., Lin, C.-Y., Kuraku, S., Hashimoto, S., Suwa, Y., Yong, L. W., Takino, K., Higashida, S., et al. (2019). Co-option of the PRDM14-CBFA2T complex from motor neurons to pluripotent cells during vertebrate evolution. Development 146,.

Leitch, H. G., Tang, W. W. C. and Surani, M. A. (2013a). Primordial germ-cell development and epigenetic reprogramming in mammals. Curr. Top. Dev. Biol. 104, 149–87.

Leitch, H. G., Mcewen, K. R., Turp, A., Encheva, V., Carroll, T., Grabole, N., Mansfield, W., Nashun, B., Knezovich, J. G., Smith, A., et al. (2013b). Naive pluripotency is associated with global DNA hypomethylation. Nat. Struct. Mol. Biol. 20, 311–316.

Ma, Z., Swigut, T., Valouev, A., Rada-Iglesias, A. and Wysocka, J. (2011). Sequence-specific regulator Prdm14 safeguards mouse ESCs from entering extraembryonic endoderm fates. Nat. Struct. Mol. Biol. 18, 120–7.

Mak, W., Nesterova, T. B., de Napoles, M., Appanah, R., Yamanaka, S., Otte, A. P. and Brockdorff, N. (2004). Reactivation of the paternal X chromosome in early mouse embryos. Science 303, 666–9.

Moindrot, B. and Brockdorff, N. (2016). RNA binding proteins implicated in Xist-mediated chromosome silencing. Semin. Cell Dev. Biol. 56, 58–70.

Nady, N., Gupta, A., Ma, Z., Swigut, T., Koide, A., Koide, S. and Wysocka, J. (2015). ETO family protein Mtgr1 mediates Prdm14 functions in stem cell maintenance and primordial germ cell formation. Elife 4, e10150.

Nakaki, F., Hayashi, K., Ohta, H., Kurimoto, K., Yabuta, Y. and Saitou, M. (2013a). Induction of mouse germ-cell fate by transcription factors in vitro. Nature 501, 222–6.

Nakaki, F., Hayashi, K., Ohta, H., Kurimoto, K., Yabuta, Y. and Saitou, M. (2013b). Induction of mouse germ-cell fate by transcription factors in vitro. Nature 501, 222–6.

Okashita, N., Kumaki, Y., Ebi, K., Nishi, M., Okamoto, Y., Nakayama, M., Hashimoto, S., Nakamura, T., Sugasawa, K., Kojima, N., et al. (2014). PRDM14 promotes active DNA demethylation through the ten-eleven translocation (TET)-mediated base excision repair pathway in embryonic stem cells. Development 141, 269–80.

Pasque, V. and Plath, K. (2015). X chromosome reactivation in reprogramming and in development. Curr. Opin. Cell Biol. 37, 75–83.

Payer, B. (2016). Developmental regulation of X-chromosome inactivation. Semin. Cell Dev. Biol. 56, 88–99.

Payer, B. (2017). Epigenetic Regulation of X-Chromosome Inactivation. In Epigenetics in Human Reproduction and Development, pp. 113–158. World Scientific Publishing.

Payer, B. and Lee, J. T. (2014). Coupling of X-Chromosome reactivation with the pluripotent stem cell state. RNA Biol. 11, 798–807.

Payer, B., Rosenberg, M., Yamaji, M., Yabuta, Y., Koyanagi-Aoi, M., Hayashi, K., Yamanaka, S., Saitou, M. and Lee, J. T. (2013). Tsix RNA and the germline factor, PRDM14, link X reactivation and stem cell reprogramming. Mol. Cell 52, 805–18.

Plath, K., Fang, J., Mlynarczyk-Evans, S. K., Cao, R., Worringer, K. A., Wang, H., de la Cruz, C. C., Otte, A. P., Panning, B. and Zhang, Y. (2003). Role of histone H3 lysine 27 methylation in X inactivation. Science 300, 131–5.

Reik, W. and Surani, M. A. (2015). Germline and Pluripotent Stem Cells. Cold Spring Harb. Perspect. Biol. 7, 1–24.

Saitou, M., Kagiwada, S. and Kurimoto, K. (2012). Epigenetic reprogramming in mouse pre-implantation development and primordial germ cells. Development 139, 15–31.

Schindelin, J., Arganda-Carreras, I., Frise, E., Kaynig, V., Longair, M., Pietzsch, T., Preibisch, S., Rueden, C., Saalfeld, S., Schmid, B., et al. (2012). Fiji: An open-source platform for biological-image analysis. Nat. Methods 9, 676–682.

Schoeftner, S., Sengupta, A. K., Kubicek, S., Mechtler, K., Spahn, L., Koseki, H., Jenuwein, T. and Wutz, A. (2006). Recruitment of PRC1 function at the initiation of X inactivation independent of PRC2 and silencing. EMBO J. 25, 3110–22.

Seisenberger, S., Andrews, S., Krueger, F., Arand, J., Walter, J., Santos, F., Popp, C., Thienpont, B., Dean, W. and Reik, W. (2012). The dynamics of genome-wide DNA methylation reprogramming in mouse primordial germ cells. Mol. Cell 48, 849–62.

Seki, Y. (2018). PRDM14 Is a Unique Epigenetic Regulator Stabilizing Transcriptional Networks for Pluripotency. Front. cell Dev. Biol. 6, 12.

Seki, Y., Hayashi, K., Itoh, K., Mizugaki, M., Saitou, M. and Matsui, Y. (2005). Extensive and orderly reprogramming of genome-wide chromatin modifications associated with specification and early development of germ cells in mice. Dev. Biol. 278, 440–58.

Seki, Y., Yamaji, M., Yabuta, Y., Sano, M., Shigeta, M., Matsui, Y., Saga, Y., Tachibana, M., Shinkai, Y. and Saitou, M. (2007). Cellular dynamics associated with the genome-wide epigenetic reprogramming in migrating primordial germ cells in mice. Development 134, 2627–38.

Silva, J. C. R., Mak, W., Zvetkova, I., Appanah, R., Nesterova, T. B., Webster, Z., Peters, A., Jenuwein, T., Otte, A. P. and Brockdorff, N. (2003). Establishment of histone h3 methylation on the inactive X chromosome requires transient recruitment of Eed-Enx1 polycomb group complexes. Dev. Cell 4, 481–495.

Sugimoto, M. and Abe, K. (2007). X chromosome reactivation initiates in nascent primordial germ cells in mice. PLoS Genet. 3, e116.

Tang, W. W. C., Dietmann, S., Irie, N., Leitch, H. G., Floros, V. I., Bradshaw, C. R., Hackett, J. A., Chinnery, P. F. and Surani, M. A. (2015). A Unique Gene Regulatory Network Resets the Human Germline Epigenome for Development. Cell 161, 1453–1467.

Tu, S., Narendra, V., Yamaji, M., Vidal, S. E., Rojas, L. A., Wang, X., Kim, S. Y., Garcia, B. A., Tuschl, T., Stadtfeld, M., et al. (2016). Co-repressor CBFA2T2 regulates pluripotency and germline development. Nature 534, 387–90.

Weber, S., Eckert, D., Nettersheim, D., Gillis, A. J. M., Schäfer, S., Kuckenberg, P., Ehlermann, J., Werling, U., Biermann, K., Looijenga, L. H. J., et al. (2010). Critical function of AP-2 gamma/TCFAP2C in mouse embryonic germ cell maintenance. Biol. Reprod. 82, 214–23.

Yamaji, M., Seki, Y., Kurimoto, K., Yabuta, Y., Yuasa, M., Shigeta, M., Yamanaka, K., Ohinata, Y. and Saitou, M. (2008). Critical function of Prdm14 for the establishment of the germ cell lineage in mice. Nat. Genet. 40, 1016–22.

Yamaji, M., Ueda, J., Hayashi, K., Ohta, H., Yabuta, Y., Kurimoto, K., Nakato, R., Yamada, Y., Shirahige, K. and Saitou, M. (2013). PRDM14 ensures naive pluripotency through dual regulation of signaling and epigenetic pathways in mouse embryonic stem cells. Cell Stem Cell 12, 368–82.

Zhao, J., Sun, B. K., Erwin, J. A., Song, J.-J. and Lee, J. T. (2008). Polycomb proteins targeted by a short repeat RNA to the mouse X chromosome. Science 322, 750–6.

