## Supplementary Information for "PRDM14 controls X-chromosomal and global epigenetic reprogramming of H3K27me3 in migrating mouse primordial germ cells"

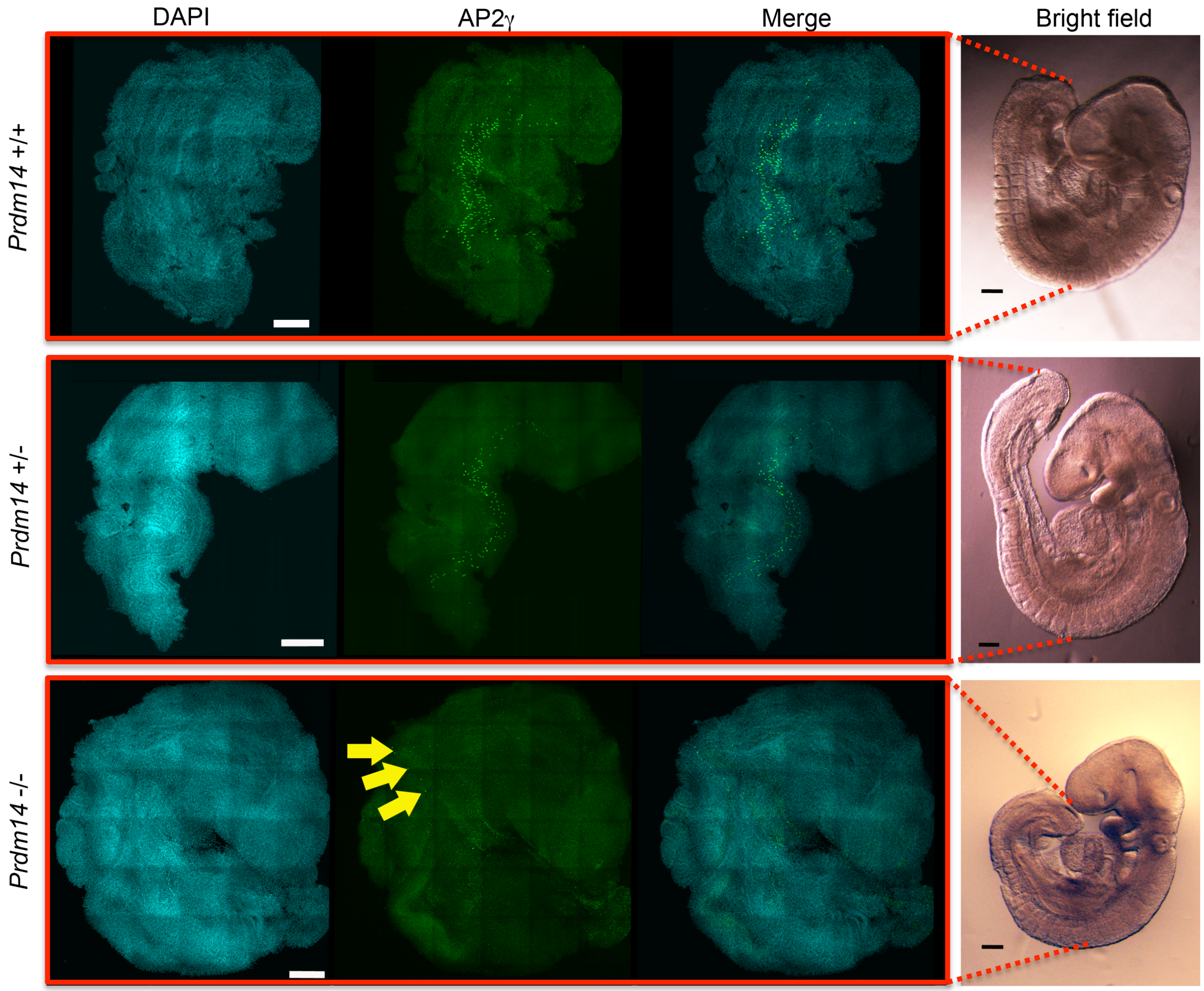


**Figure S1**

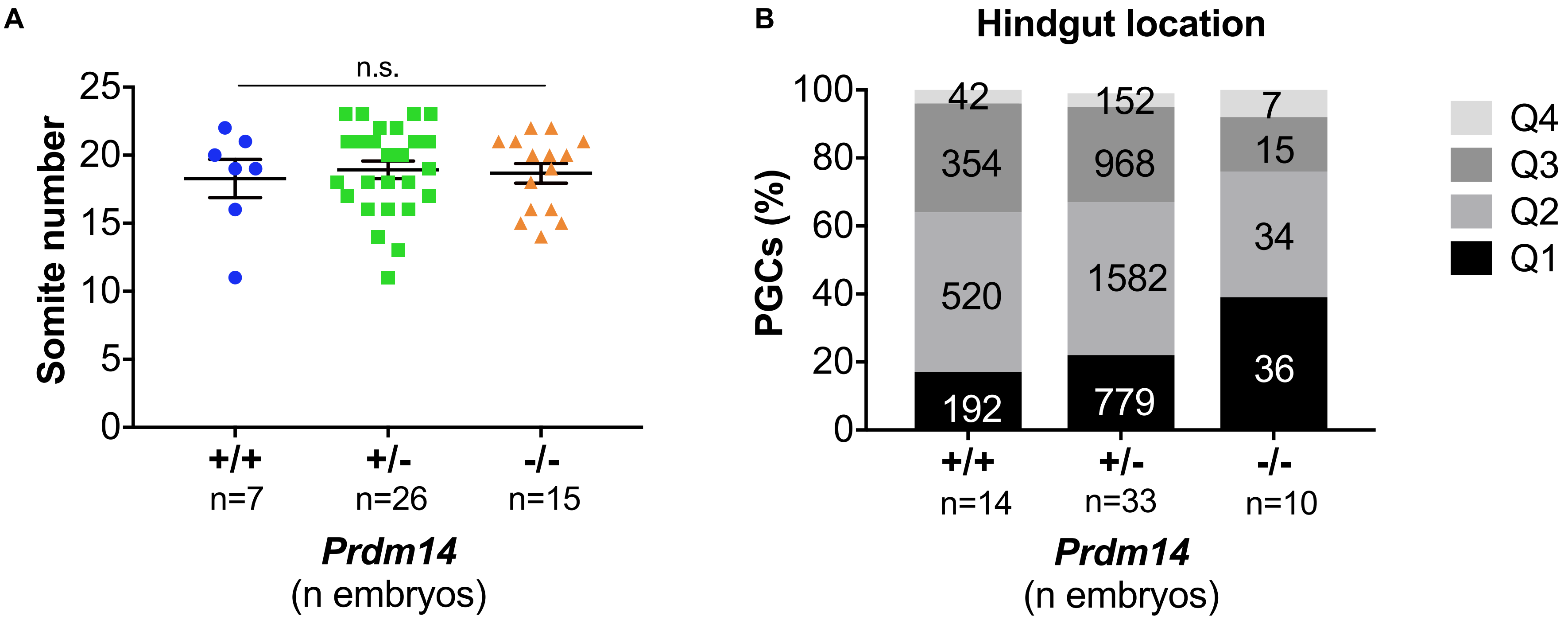
A
n.s.
Somite number
25
20
15
10
5
0
+/+
n=7
+/-
n=26
-/-
n=15
Prdm14
(n embryos)
B
Hindgut location
PGCs (%)
100
80
60
40
20
0
42
354
520
192
152
968
1582
779
7
15
34
36
Q4
Q3
Q2
Q1
+/+
n=14
+/-
n=33
-/-
n=10
Prdm14
(n embryos)

**Figure S2**

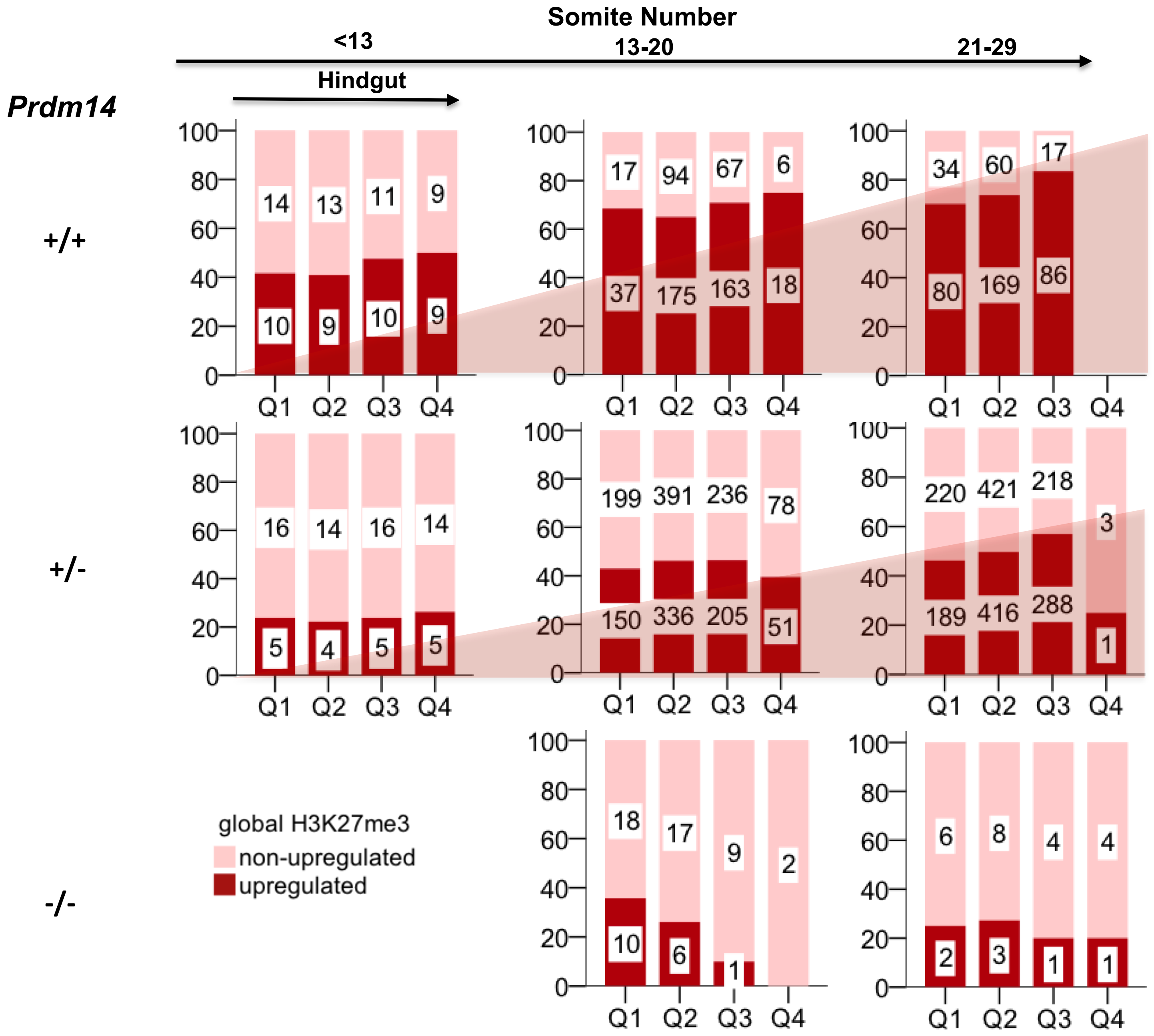

Somite Number
<13
13-20
21-29
Hindgut
Prdm14
+/+
+/-
-/-
100
80
60
40
20
0
Q1
Q2
Q3
Q4
14
13
11
9
10
9
10
9
17
94
67
6
37
175
163
18
34
60
17
80
169
86
16
14
16
14
5
4
5
5
199
391
236
78
150
336
205
51
220
421
218
3
189
416
288
1
18
17
9
2
10
6
1
6
8
4
4
2
3
1
1
global H3K27me3
non-upregulated
upregulated


Figure S3

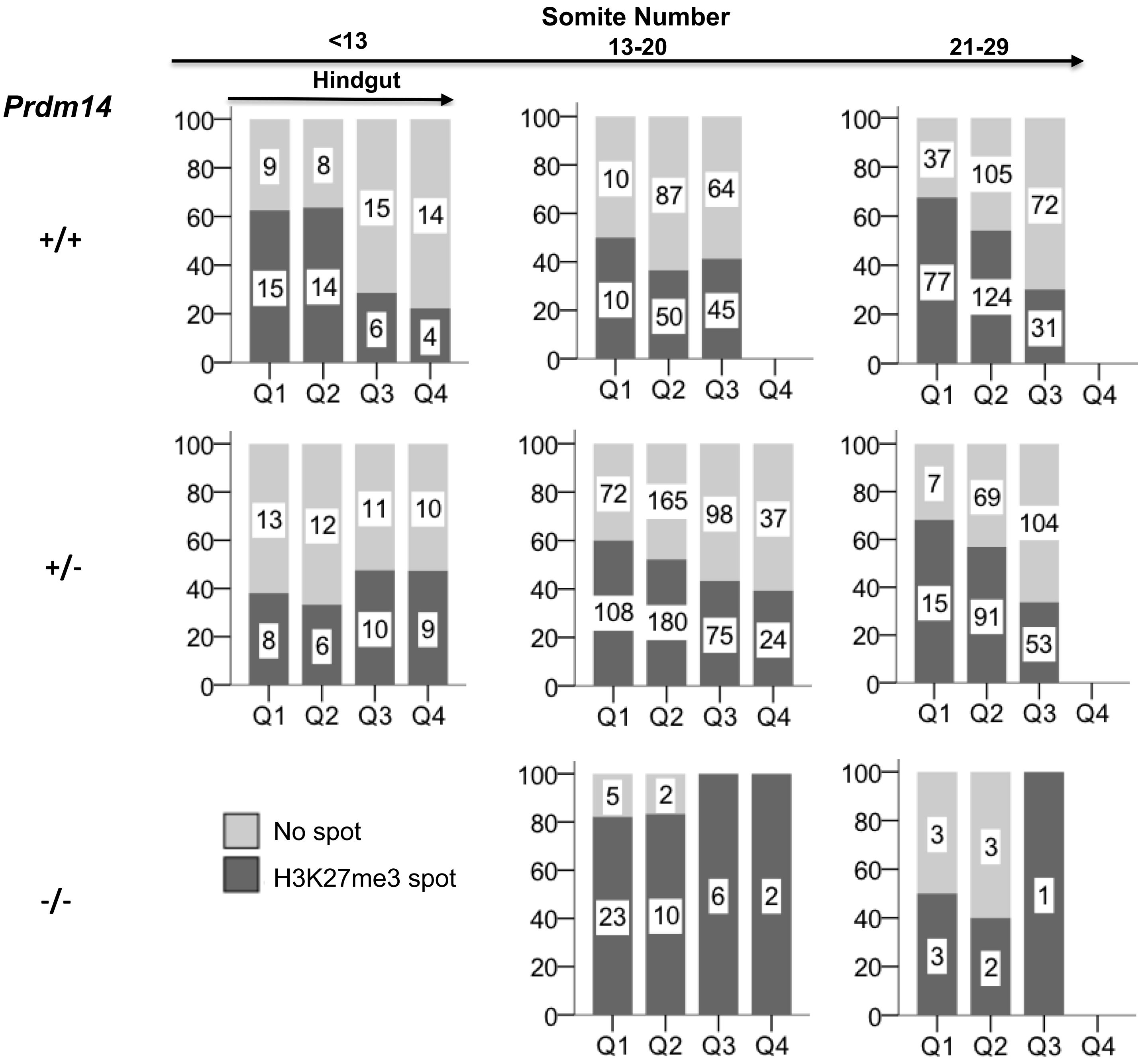
Somite Number
<13
13-20
21-29
Hindgut
Prdm14
+/+
+/-
-/-
9
8
15
14
15
14
6
4
10
87
64
10
50
45
37
105
72
77
124
31
13
12
11
10
8
6
10
9
72
165
98
37
108
180
75
24
7
69
104
15
91
53
5
2
23
10
6
2
3
3
3
2
1
Q1
Q2
Q3
Q4
No spot
H3K27me3 spot

Figure S4

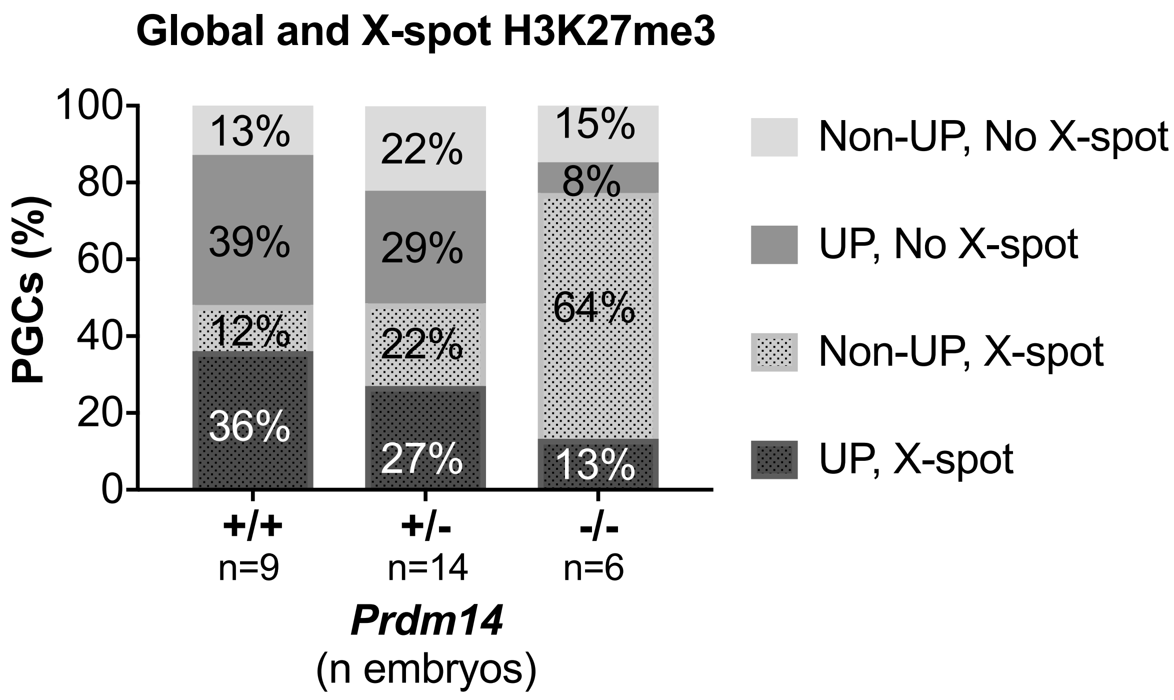

Global and X-spot H3K27me3
PGCs (%)
100
80
60
40
20
0
13%
39%
12%
36%
22%
29%
22%
27%
15%
8%
64%
13%
+/+
n=9
+/-
n=14
-/-
n=6
Prdm14
(n embryos)
Non-UP, No X-spot
UP, No X-spot
Non-UP, X-spot
UP, X-spot


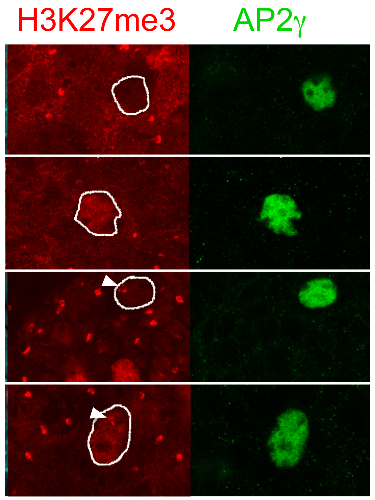

H3K27me3
AP2γ


**Figure S5**

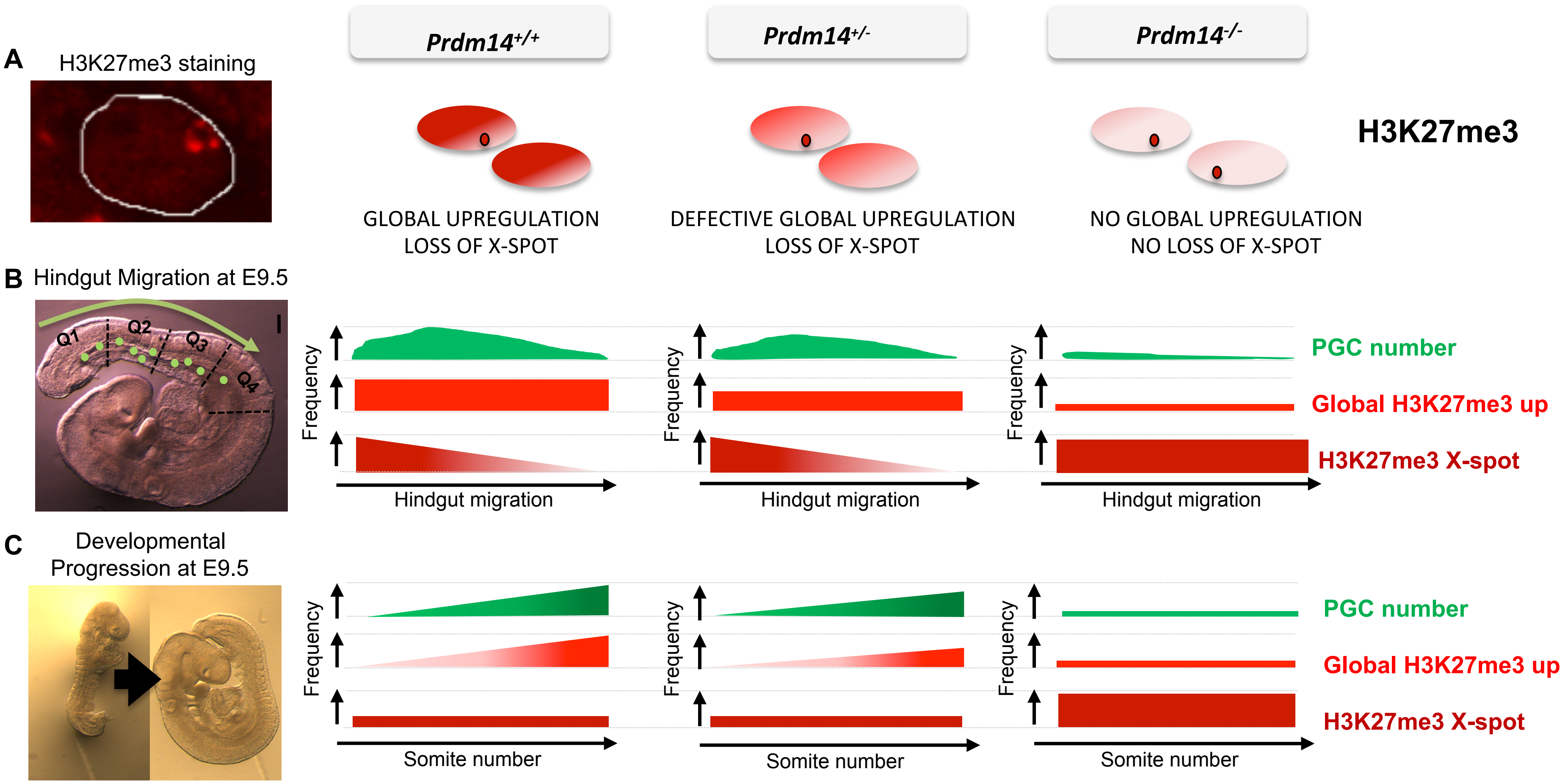

A
H3K27me3 staining
Prdm14+/+
Prdm14+/-
Prdm14-/-
H3K27me3
GLOBAL UPREGULATION
LOSS OF X-SPOT
DEFECTIVE GLOBAL UPREGULATION
LOSS OF X-SPOT
NO GLOBAL UPREGULATION
NO LOSS OF X-SPOT
B
Hindgut Migration at E9.5
Q1
Q2
Q3
Q4
Frequency
Hindgut migration
PGC number
Global H3K27me3 up
H3K27me3 X-spot
C
Developmental
Progression at E9.5
Frequency
Somite number
PGC number
Global H3K27me3 up
H3K27me3 X-spot


**Figure S6**

**SUPPLEMENTARY INFORMATION**

**Supplementary Figure Legends**

**Figure S1 (related to Fig. 1). Migrating PGCs in E9.5 mouse embryos of different *Prdm14* genotypes.** Representative immunofluorescence images of hindgut regions (left) and corresponding bright field images of E9.5 embryos (18-19 somites, right) after whole mount antibody staining. Migrating PGCs express AP2γ. Arrows indicate four PGCs in a *Prdm14*$^{-/-}$ hindgut. Scale bars: 250 μm.

**Figure S2 (related to Fig. 1). *Prdm14* genotype, developmental progression and PGC distribution during migration.** (A) The *Prdm14* genotype does not affect developmental progression in E9.5 embryos (Kruskal-Wallis test, p=0.853). Somite numbers per embryo are plotted as marker of developmental progression versus *Prdm14* genotype. Mean somite number and standard error (error bars) are given for each genotype. (B) Distribution of PGCs along the hindgut across *Prdm14* genotypes. Labels in each category indicate numbers of PGCs counted.

**Figure S3 (related to Fig. 2). Summary of global H3K27me3 levels in PGCs across somite number, migration progression and *Prdm14* genotypes.** The pink triangles symbolize the trend of increasing frequencies of PGCs with high global H3K27me3 levels with developmental progression (increasing somite number) in *Prdm14*$^{+/+}$ and *Prdm14*$^{+/-}$ (to a lesser degree than in *Prdm14*$^{+/+}$), but not in *Prdm14*$^{-/-}$ embryos.

**Figure S4 (related to Fig. 3). Summary of the removal of X-chromosomal H3K27me3 accumulation (spot) in PGCs across somite number, migration progression and *Prdm14* genotypes.** A trend in removal of the H3K27me3 spot can be observed with progressing migration of PGCs in *Prdm14*$^{+/+}$ and *Prdm14*$^{+/-}$, but not in *Prdm14*$^{-/-}$ embryos.

**Figure S5 (related to Fig. 4). Distribution of PGCs with different global and X-chromosomal H3K27me3 status dependent on *Prdm14* genotype.** The bar chart shows the frequency of AP2γ-positive PGCs of the different H3K27me3 categories depicted in the immunostaining images on the right, in relation to *Prdm14* genotype. UP, global upregulated H3K27me3 staining in relation to surrounding somatic cells; X-spot, X-chromosomal H3K27me3 accumulation (white arrowheads).

**Figure S6. Summary of H3K27me3 dynamics in PGCs.** (A) Schematic of H3K27me3 staining in representative female PGC nuclei of different *Prdm14* genotypes. (B) PGC numbers and H3K27me3 changes during PGC migration along the hindgut at E9.5. PGCs are most frequently seen in quadrant 2 (Q2) and their number is strongly reduced in *Prdm14*$^{-/-}$ embryos. Global H3K27me3 levels are upregulated in most PGCs in *Prdm14*$^{+/+}$ embryos throughout the hindgut, while it is less frequent in *Prdm14*$^{+/-}$ and rarely seen in *Prdm14*$^{-/-}$ embryos. The accumulation of H3K27me3 on the inactive X-chromosome (X-spot) is progressively lost during migration in both *Prdm14*$^{+/+}$ and *Prdm14*$^{+/-}$ PGCs, but mostly maintained in *Prdm14*$^{-/-}$ PGCs. (C) PGC numbers and H3K27me3 changes in E9.5 embryos with different developmental progression (somite number). PGC numbers and global H3K27me3 levels increase with developmental progression in *Prdm14*$^{+/+}$ and *Prdm14*$^{+/-}$ (reduced upregulation), but not in *Prdm14*$^{-/-}$ embryos. The X-chromosomal H3K27me3 spot is lost both in *Prdm14*$^{+/+}$ and *Prdm14*$^{+/-}$ PGCs independently of developmental progression, while *Prdm14*$^{-/-}$ PGCs maintain the X-spot at all stages.

### Supplementary Table S1

| Genotype | N Embryos (male/female) | Somite number (mean ± SD) | PGC number (mean ± SD) | % Global H3K27me3 UP (mean ± SD) | % loss of H3K27me3 X-spot (mean ± SD) |
|---|---|---|---|---|---|
| *Prdm14* +/+ | 14 (5/9) | 18.29 ± 3.73[a] | 153.43 ± 44.01[a] | 76.26 ± 17.38[a] | 52.91 ± 8.58[a] |
| *Prdm14* +/- | 33 (19/14) | 18.92 ± 3.27[a] | 134.85 ± 37.89[a] | 52.30 ± 18.57[b] | 50.61 ± 7.51[a] |
| *Prdm14* -/- | 17 (7/10) | 18.67 ± 2.77[a] | 7.53 ± 7.41[b] | 21.03 ± 12.94[c] | 21.16 ± 17.41[b] |

[a,b,c] Values with different superscripts differ significantly from each other within the same column in the Kruskal-Wallis test ($P<0.05$).
